# Transformation of European ash (*Fraxinus excelsior* L.) callus as a starting point for understanding the molecular basis of ash dieback

**DOI:** 10.1101/2020.12.03.409425

**Authors:** Anna Hebda, Aleksandra Liszka, Piotr Zgłobicki, Katarzyna Nawrot-Chorabik, Jan J. Lyczakowski

**Author notes:** Corresponding author: Jan J. Lyczakowski. **Key message:** We report the first protocol for genetic transformation of European ash (*Fraxinus excelsior* L.) callus tissue and selection conditions allowing for generation of transgenic callus lines.

## Abstract

Plant genetic engineering requires transfer of genetic material into plant cells, a process that is frequently challenging for tree species. Here we report first successful genetic transformation of European ash (*Fraxinus excelsior*). The protocol relies on the use of *Agrobacterium tumefaciens* to transform callus tissue derived from embryos of *F. excelsior*. In our experiments, we use the β-glucuronidase (GUS) reporter system to demonstrate transformation of ash callus tissue. Moreover, we describe an antibiotic-resistance based selection process enabling formation of stable transgenic callus lines. Since ash dieback threatens the long-term stability of many native *F. excelsior* populations, we hope that the transformation techniques described in this manuscript will facilitate rapid progress in uncovering the molecular basis of the disease and mechanisms used by trees to resist it.

## Introduction

European ash (*Fraxinus excelsior* L.) is a deciduous woody hardwood species with a broad geographical distribution and high-quality timber. These properties make *F. excelsior* both ecologically and economically important (Dobrowolska et al., 2011). The population of *F. excelsior* is currently facing a risk of collapse, mainly due to ash dieback, a disease caused by pathogenic fungus *Hymenoscyphus fraxineus* (Kowalski, 2006). The mortality of *F. excelsior* due to the dieback can reach as much as 85% in certain populations, but interestingly some individuals are largely resistant to the disease (Coker et al., 2019). The progress in understanding ash dieback has been greatly enhanced by solving *H*. *fraxineus* (McMullan et al., 2018) and *F. excelsior* genomes (Sollars et al., 2017) and by large sequencing projects of healthy and diseased trees (Stocks et al., 2019). However, rapid advances in uncovering the biological basis of the disease remain somehow impeded, partially due to lack of genetic engineering tools for *F. excelsior.* Previous publications reported transformation of other ash species, such as *Fraxinus pensylvanica* (Du and Pijut, 2009), *Fraxinus americana* (Palla and Pijut, 2015) and *Fraxinus profunda* (Stevens and Pijut, 2014). Here we describe the first protocol for transformation of *F. excelsior* callus tissue. In addition, we outline the selection conditions, allowing for generation of transgenic callus lines. Since recalcitrance to transformation is one of the main challenges in tree genetic engineering (Song et al., 2019), we believe that our protocol will facilitate progress in using reverse genetics approaches to study *F. excelsior* and the *H. fraxineus* pathogenicity.

## Results

In order to establish a transformation protocol for *F. excelsior* callus, we decided to employ *Agrobacterium tumefaciens* bearing a plasmid encoding GUS and a kanamycin resistance selectable marker. Promoter 35S (p35S), used to drive GUS expression, is active in a range of plant species and together with GUS reporter it was successfully used in other *Fraxinus* species (Du and Pijut, 2009, Palla and Pijut, 2015, Stevens and Pijut, 2014). To assess if our protocol allows for transformation of *F. excelsior* cells, we incubated the callus with the X-Gluc substrate solution ten days after initial infiltration with *A. tumefaciens.* As a control, we analysed callus which was subjected to same procedures but infiltrated in a suspension of *A. tumefaciens* free of the p35S:GUS plasmid. Our analysis indicated that some of the callus fragments incubated with the p35S:GUS plasmid bearing *A. tumefaciens* developed blue colouration, which is likely caused by GUS activity (Figure 1A). None of the control callus fragments developed blue colouration. In our preliminary experiments we tested a range of *A. tumefaciens* strains, including AGL-1, GV3101 and C58. Importantly, we were only able to observe blue colouration on the callus when AGL-1 *A. tumefaciens* was used.

**Figure 1.**
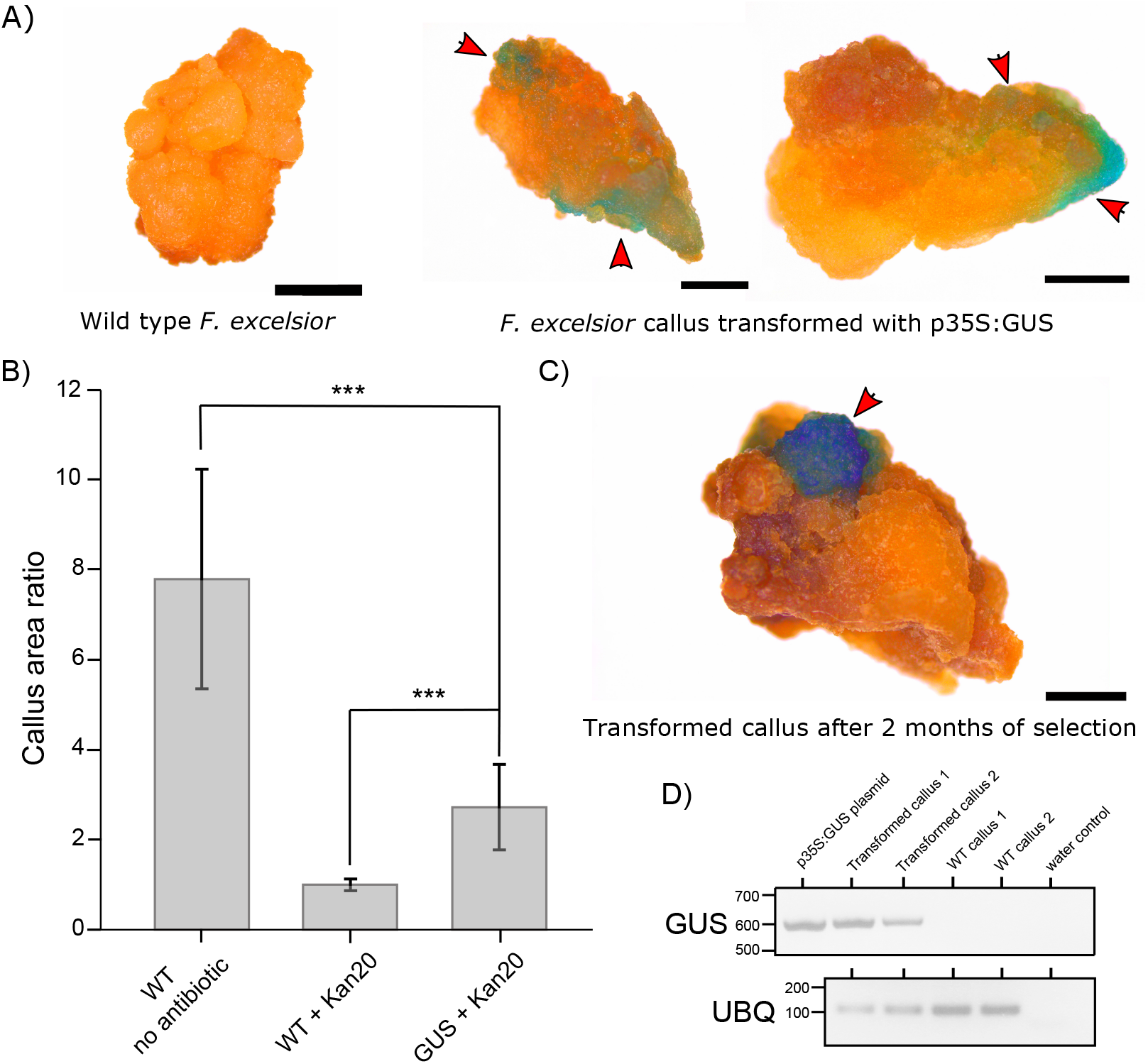
Transformation of *F. excelsior* callus. **A)** Control (left) and transformed (right) callus fragments stained for GUS activity 10 days after incubation with AGL-1 *A. tumefaciens*. **B)** Comparison of area increase for WT callus grown on medium without (WT no antibiotic) and with 20 ug/mL kanamycin (WT + Kan20) and for callus transformed with the pORE-R1:prom35S:GUS construct (GUS + Kan20). **C)** Transformed callus fragment stained for GUS activity eight weeks after incubation with AGL-1 *A. tumefaciens*. **D)** RT-PCR results for *GUS* (top) and *UBIQUITIN* (*UBQ*, bottom) amplified for two transformed and WT calli. For GUS a purified pORE-R1:prom35S:GUS was used as a template in a positive control reaction. Water was used as a template in negative control reactions. Size bars in A) and C) correspond to 1 mm, red arrows indicate zones with GUS activity. Student’s T-test was used to analyse statistical significance in B), *** denotes p ≤ 0.001.

To enable generation of stable genetically modified *F. excelsior* lines, it is necessary to develop a selection process for the transformed cells. In our experiments, we used resistance to kanamycin to facilitate the selection process. Firstly, we analysed the growth of WT *F. excelsior* callus on solid medium supplemented with a range of kanamycin concentrations (SI Figure 2). This experiment enabled us to establish that supplementation of medium with 20 ug/mL kanamycin is sufficient to completely inhibit WT callus growth (SI Figure 2). To quantify the efficiency of the selection process we measured the area change for WT calli grown on medium with and without the addition of kanamycin (Figure 1B). Over an eight-week period, the WT callus increased its area by around eight times on medium without kanamycin (Figure 1B). Over the same period, callus fragments grown on medium with 20 ug/mL kanamycin did not increase their area. This indicates that supplementation of medium with 20 ug/mL of kanamycin is enough to effectively stop the growth of WT callus.

Having established suitable selection conditions, we measured area increase for calli transformed with our construct, which, in addition to encoding the GUS protein, confers resistance to kanamycin. On average, the transformed calli increased their area more than two times over an eight-week period (Figure 1B). This confirms that cell division had occurred in the transgenic callus and indicates that transformed calli were resistant to kanamycin, what was likely conferred by the inserted transgene. To provide further evidence for stable transformation of the callus we performed GUS staining on the callus fragments eight weeks after transformation. We observed GUS staining (Figure 1C, SI Figure 3) for parts of the transformed fragment, which may be attributed to newly formed cell material.

Since it is possible that the GUS protein remains stable in cells only transiently transformed with the construct, we wanted to validate that the *GUS* gene is expressed in callus grown on medium supplemented with kanamycin over an eight-week period. To this end, we performed RT-PCR for *GUS* on WT and transformed calli (Figure 1D). Amplicon was generated from cDNA of transgenic callus and no amplification was observed for WT. As a housekeeping control we amplified a 3’ end of *UBIQUITIN*. In this reaction, we observed an amplicon when using cDNA from both WT and the transgenic callus. This experiment indicates constant *GUS* mRNA production in transgenic callus and provides further evidence for a stable integration of the transgene into *F. excelsior* genome.

## Discussion and conclusion

In this manuscript we describe a protocol for transformation of callus tissue of *F. excelsior*. In our experiments we use *GUS* as a reporter gene and apply kanamycin selection and RT-PCR to provide further evidence for stable integration of the transgene into *F. excelsior* genome. Our protocol shares many similarities with previous experiments describing transformation of other ash species (Du and Pijut, 2009, Palla and Pijut, 2015, Stevens and Pijut, 2014). For example, like ours, published protocols apply both sonication and infiltration to facilitate *A. tumefaciens* transfer into plant cells. Moreover, in line with our results, for most investigated species, with a notable exception of *F. americana* (Palla and Pijut, 2015), supplementation of medium with 20 ug/mL kanamycin is sufficient to select transformed cells. A key difference between our work and previously published protocols lies in the type of plant material used for transformation. While other groups transformed hypocotyls, which were then used for regeneration of transgenic callus, we transformed callus tissue directly.

In our experiments, it is important to distinguish transient and stable transformation events. Similarly to results observed for hypocotyls of *F. pensylvanica* (Du and Pijut, 2009), some of our transformed callus fragments produced blue pigment, associated with GUS activity, in multiple spots when stained ten days after infiltration with *A. tumefaciens*. Importantly, after eight weeks of incubation on the selection medium, a smaller number of zones with GUS activity was observed on calli (Figure 1A and C). This suggests that initially multiple cells can be transformed in one callus fragment but most of the initial transient *GUS* expression is likely to be lost over time. This prevents accurate quantification of transformation efficiency obtained in our experiments. Importantly, stable maintenance of the transgene in only some of the transformed cells also might explain why the transgenic calli increased their area two times over the eight weeks of selection, while, over the same time, the WT callus grew more than eight times on medium without kanamycin. Alternatively, this discrepancy in growth rate might be associated with metabolic burden of transgene expression by transformed cells.

In summary, our work demonstrates effective transformation of *F. excelsior* callus. We hope that our protocol may find application in generation of transgenic trees which can be used in studies aimed at understanding the biological basis of ash dieback.

## Materials and Methods

### Plant material used and culturing conditions

Experiments were performed on *in vitro* grown callus tissue. Zygotic embryos isolated from mature *F. excelsior* seeds were used as initial explants. Allogamic seeds were obtained from Kostrzyca Forest Gene Bank and originated from Jarocin district of the General Directorate of the State Forests located in western Poland (51°58’3.863“N, 17°29’53.772”E, 120 masl). Callus was derived and grown on a medium described in a separate patent application (Nawrot-Chorabik and Latowski, 2020). Calli of between 4 and 5 cm in diameter were cut into sections of up to 0.5 cm in diameter, which were used for transformation.

### Callus transformation and selection

A detailed description of the transformation protocol is provided in SI Figure 1. In brief, callus fragments were placed in a culture of AGL-1 *A. tumefaciens* bearing the pORE-R1:prom35S:GUS plasmid (Coutu et al., 2007) and sonicated. Following sonification, bacteria were vacuum infiltrated into callus fragments. Infiltrated calli were placed on solid medium and incubated in the dark for 48 hours. After incubation, callus fragments were washed in water supplemented with 100 ug/mL Timentin and placed on solid medium amended with Kanamycin (variable concentrations used) and 100 ug/mL Timentin.

### GUS staining and imaging

Solution (100 mg/dm^3^) of X-Gluc (ThermoFisher Scientific) in 100 mM pH 7.0 sodium phosphate buffer was used for GUS staining. Solution was vacuum infiltrated into callus fragments and immersed calli were left overnight at 37°C to allow for GUS catalysed reaction. Chlorophyll removal was performed with 96% ethanol. Callus images were obtained with a stereomicroscope (Leica S9i).

### RT-PCR analysis of gene expression

Total RNA was extracted from callus fragments grown on selective medium for eight weeks using a commercial kit (Sigma, Spectrum™ Plant Total RNA). First strand cDNA was synthesised using polyA primers and RevertAid First Strand cDNA synthesis kit (ThermoFisher Scientific). For each RT-PCR reaction, 500 ng of cDNA was used. Primer sequences and PCR conditions used to amplify *GUS* and *UBIQUITIN* (*UBQ*) fragments are provided in SI Table 1.

## Author contributions

AH and JJL designed the study, performed callus transformation and RT-PCR analysis, AL supported callus maintenance and selection, PZ prepared the genetic construct used for transformation, KNCH designed the study, derived callus lines used for transformation and prepared growth media, JJL, AH and KNCH wrote the manuscript.

## Acknowledgements

In memory of Dr Damian Ryszawy, who trained us in the use of a stereomicroscope applied in this project. This work was supported by a grant from the National Science Centre, Poland awarded to JJL as part of the SONATINA 3 programme (project number 2019/32/C/NZ3/00392).

## Supporting information figure and table list

SI Figure 1 – Detailed description of the transformation protocol.

SI Figure 2 – Selection conditions evaluated for *F. excelsior* callus.

SI Figure 3 – Further images of GUS activity in callus eight weeks after transformation.

SI Table 1 – Primer sequences and RT-PCR conditions used.

## List of abbreviations

GUS: β-glucuronidase enzyme
p35S: promoter 35S
X-Gluc: 5-bromo-4-chloro-3-indolyl-beta-D-glucuronic acid
Kan: Kanamycin
RT-PCR: Reverse transcription polymerase chain reaction
UBQ: Ubiquitin

## References

Coker, T. L. R., RozsypÁlek, J., Edwards, A., Harwood, T. P., Butfoy, L. & Buggs, R. J. A. 2019. Estimating mortality rates of European ash (F*raxinus excelsior)* under the ash dieback (H*ymenoscyphus fraxineus)* epidemic. Plants, People, Planet, 1,48–58.

Coutu, C., Brandle, J., Brown, D., Brown, K., Miki, B., Simmonds, J. & Hegedus, D. D. 2007. pORE: a modular binary vector series suited for both monocot and dicot plant transformation. Transgenic Research, 16,771–781.

Dobrowolska, D., Hein, S., Oosterbaan, A., Wagner, S., Clark, J. & Skovsgaard, J. P. 2011. A review of European ash (Fraxinus excelsior L.): implications for silviculture. Forestry, 84,133–148.

Du, N. X. & Pijut, P. M. 2009. Agrobacterium-mediated transformation of Fraxinus pennsylvanica hypocotyls and plant regeneration. Plant Cell Reports, 28,915–923.

Kowalski, T. 2006. Chalara fraxinea sp nov associated with dieback of ash (Fraxinus excelsior) in Poland. Forest Pathology, 36,264–270.

Mcmullan, M., Rafiqi, M., Kaithakottil, G., Clavijo, B. J., Bilham, L., Orton, E., Percival-Alwyn, L., Ward, B., Edwards, A., Saunders, D. G. O., Accinelli, G. G., Wright, J., Verweij, W., Koutsovoulos, G., Yoshida, K., Hosoya, T., Williamson, L., Jennings, P., Ioos, R., Husson, C., Hietala, A. M., Vivian-SMITH, A., Solheim, H., Maclean, D., Fosker, C., Hall, N., Brown, J. K. M., Swarbreck, D., Blaxter, M., Downie, J. A. & Clark, M. D. 2018. The ash dieback invasion of Europe was founded by two genetically divergent individuals. Nature Ecology & Evolution, 2,1000-+.

Nawrot-Chorabik, K. & Latowski, D. 2020. Sposób pozyskiwania sadzonek jesionu wyniosłego (Fraxinus excelsior L.) oraz pożywki nadające się do stosowania w tym sposobie. P.433288 WIPO ST 10/C PL433288.

Palla, K. J. & Pijut, P. M. 2015. Agrobacterium-mediated genetic transformation of Fraxinus americana hypocotyls. Plant Cell Tissue and Organ Culture, 120,631–641.

Sollars, E. S. A., Harper, A. L., Kelly, L. J., Sambles, C. M., Ramirez-Gonzalez, R. H., Swarbreck, D., Kaithakottil, G., Cooper, E. D., Uauy, C., Havlickova, L., Worswick, G., Studholme, D. J., Zohren, J., Salmon, D. L., Clavijo, B. J., Li, Y., He, Z. S., Fellgett, A., Mckinney, L. V., Nielsen, L. R., Douglas, G. C., Kjaer, E. D., Downie, J. A., Boshier, D., Lee, S., Clark, J., Grant, M., Bancroft, I., Caccamo, M. & Buggs, R. J. A. 2017. Genome sequence and genetic diversity of European ash trees. Nature, 541,212-+.

Song, G. Q., Prieto, H. & Orbovic, V. 2019. Agrobacterium-Mediated Transformation of Tree Fruit Crops: Methods, Progress, and Challenges. Frontiers in Plant Science, 10.

Stevens, M. E. & Pijut, P. M. 2014. Agrobacterium-mediated genetic transformation and plant regeneration of the hardwood tree species Fraxinus profunda. Plant Cell Reports, 33,861–870.

Stocks, J. J., Metheringham, C. L., Plumb, W. J., Lee, S. J., Kelly, L. J., Nichols, R. A. & Buggs, R. J. A. 2019. Genomic basis of European ash tree resistance to ash dieback fungus. Nature Ecology & Evolution, 3,1686-+.

