## Supporting_Information_File for "Transformation of European ash (*Fraxinus excelsior* L.) callus as a starting point for understanding the molecular basis of ash dieback"

**SI Table 1. Primer sequences used in this study.** RT-PCR was performed with 10 uM forward and reverse primer concentrations using PCR-Mix Plus (A&A Biotechnology). For each reaction initial melting was performed for 2 minutes at 95 °C. PCR was performed for 35 cycles with 30 seconds melting (95 °C), 45 seconds annealing (60 °C) and 90 seconds extension (72 °C). Final extension was performed for 5 minutes.

| Primer name and target | Primer sequence (5’-3’) |
|---|---|
| Fraxinus_excelsior_UBIQUITIN_For | GACCAGCAGCGATTGATCTTT |
| Fraxinus_excelsior_UBIQUITIN_Rev | GAGGACAAGATGGAGGGTAGAC |
| GUS_For | CAACGAACTGAACTGGCAGA |
| GUS_Rev | AGAGGTTAAAGCCGACAGCA |

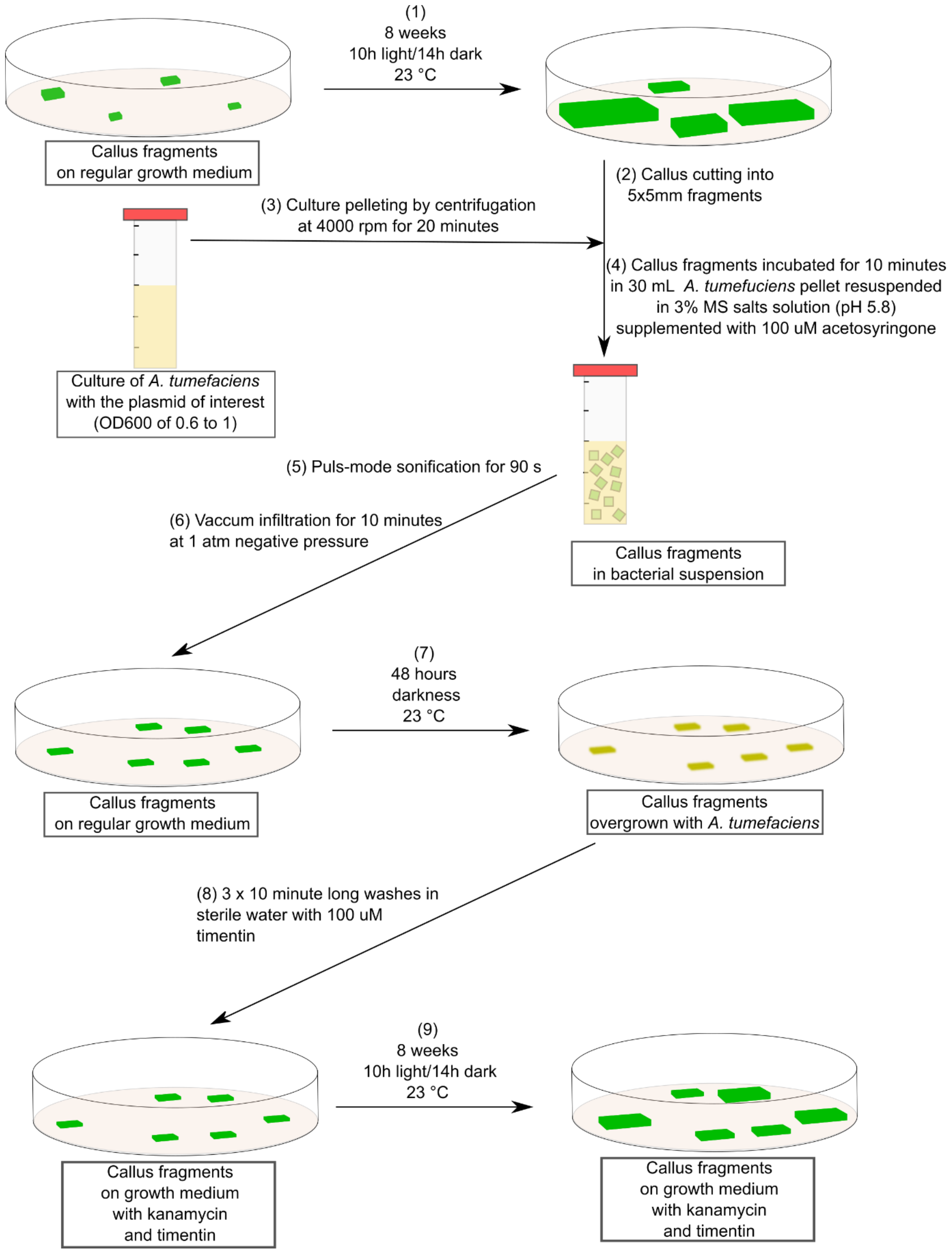


**SI Figure 1. A Detailed transformation protocol for *Fraxinus excelsior*.** Transformation steps are denoted with numbers (1) to (9). Material used for each step is described in text boxes underneath the drawing.

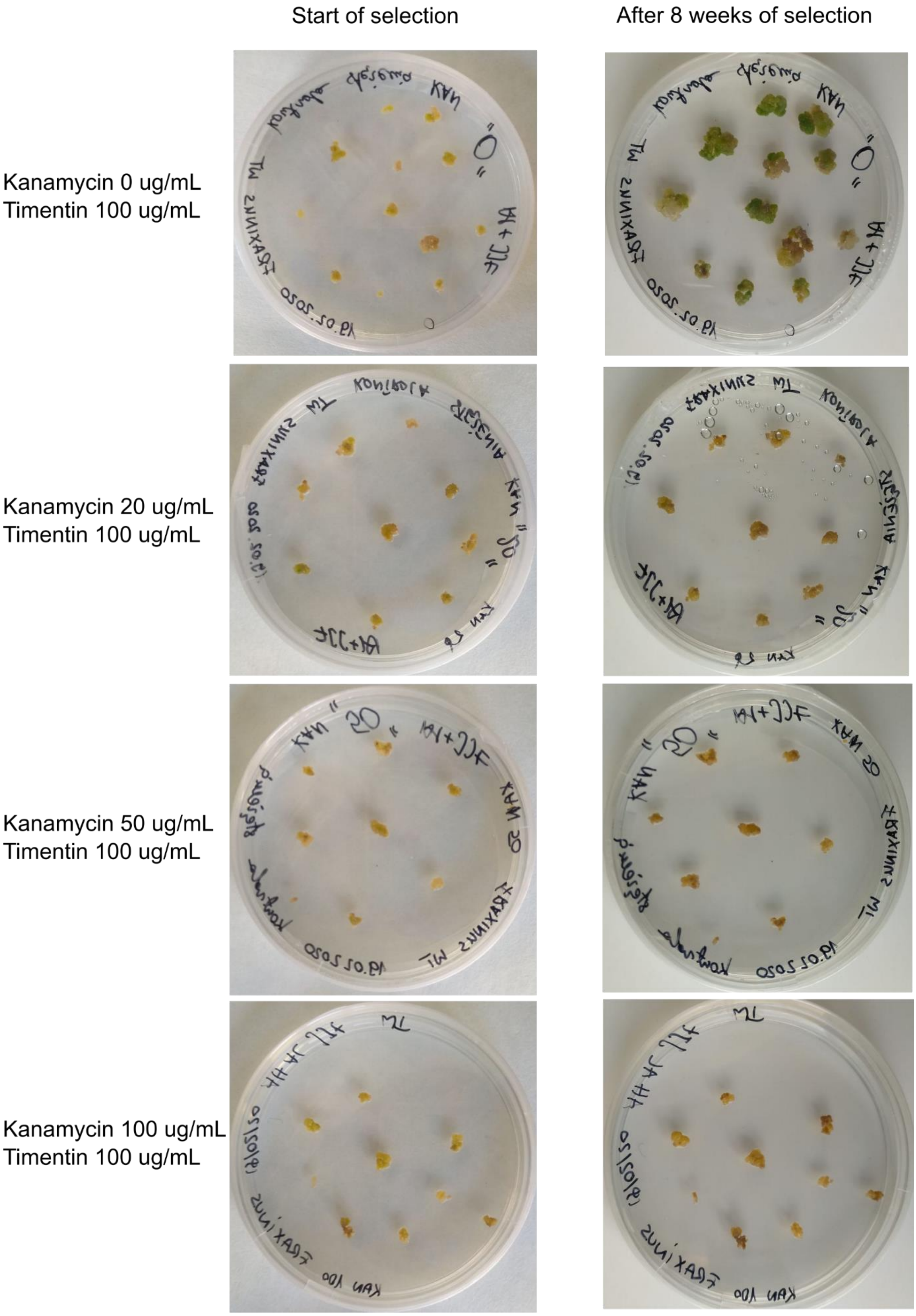


**SI Figure 2. Selection conditions evaluated for *F. excelsior* callus.** Kanamycin and timentin concentrations are provided for each image.

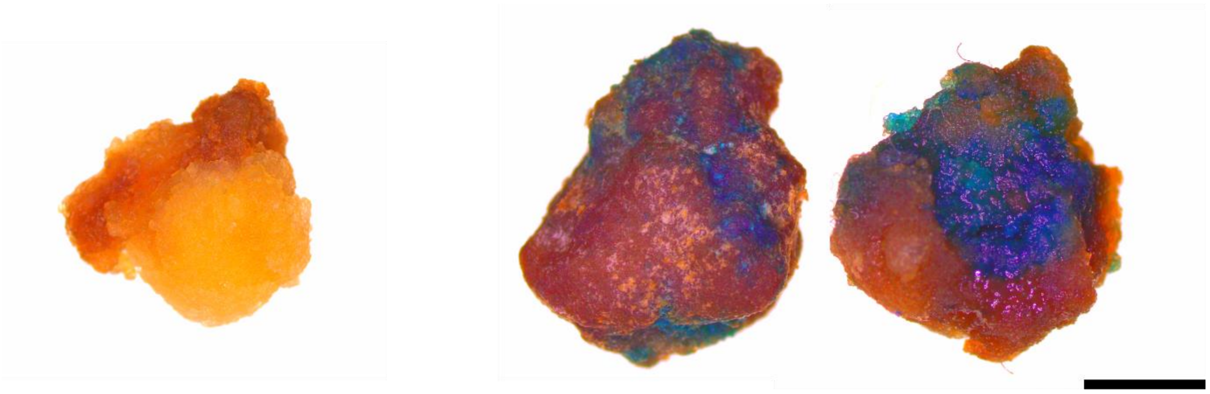

Wild type *F. excelsior* callus p35S:GUS *F. excelsior* callus

**SI Figure 3. Further images of GUS activity in callus eight weeks after transformation.** Black size bar corresponds to 1 mm.
